# Curiosity is associated with enhanced tonic firing in dorsal anterior cingulate cortex

**DOI:** 10.1101/2020.05.25.115139

**Authors:** Maya Zhe Wang, Benjamin Yost Hayden

## Abstract

Disparity between current and desired information, known as information gap, is an important driver of information-seeking and curiosity. To gain insight into its neural basis, we recorded responses of single neurons in dorsal anterior cingulate cortex (dACC) while rhesus macaques performed a task that induces and quantifies demand for information. We find that enhanced firing rates in dACC before the start of a trial predict a stronger bias towards information-seeking choices. Following choices of uninformative options, firing rates are tonically enhanced until information is delivered. The level of enhancement observed is correlated on a trial-by-trial basis with the value assigned to the prospective information. Finally, variation in this tone is positively correlated with receptiveness to new information, as inferred by preference changes on subsequent trials. These patterns are not observed in a complementary dataset collected in orbitofrontal cortex (OFC), suggesting these effects reflect at least somewhat anatomically localized processing.

## INTRODUCTION

Ignorance is not always bliss. A decision-maker who is uncertain about the outcomes of their potential actions and choices may have a desire to probe the environment for information that can provide the missing knowledge. Indeed, decision-makers may gain utility from doing so, even if the information is neutral or bad (Kidd and Hayden, 2016; White et al., 2019). This fact has motivated scholars to propose that curiosity is motivated in part by an information gap, ego dystonic discrepancy between current and desired information (Golman & Loewenstein, 2015; Gottlieb et al., 2013; Kang et al., 2009; Loewenstein, 1994). In this view, lack of information is a special drive state that can be sated by obtaining information. The information gap is the central theoretical structure linking curiosity to psychology and ultimately to neuroscience (Golman & Loewenstein, 2018; Gottlieb & Oudeyer, 2018; Kidd & Hayden, 2016; Marvin & Shohamy, 2016; van Lieshout et al., 2018).

Despite its value in motivating psychological hypotheses, the neuronal basis of the information gap remains to be identified (Cervera et al., 2020). We hypothesized that the brain computes and represents the demand for information within a circumscribed circuit. Several factors motivated us to hypothesize that the dorsal anterior cingulate cortex (dACC) would be one such region. The dACC is associated with monitoring both cognitive and visceral (i.e. basal drive state) variables (Heilbronner & Hayden, 2016a; Morecraft & Van Hoesen, 1998). At least one study has linked activity in dACC to curiosity (Jepma et al., 2012). Neurons in dACC also track - and drive demand for - counterfactual information, suggesting the region may monitor current information gap, and drive information-seeking decisions (Hayden, et al., 2009). Moreover, enhanced hemodynamic activity in this region is associated with enhanced control, with specification of control, and with exploratory processes in foraging, which have some heuristically similarity to information-seeking (Kolling et al., 2012; Shenhav et al., 2013; Shenhav et al., 2017; Smith et al., 2019; Heilbronner and Hayden, 2016). Finally, activity in this region is directly associated with information-seeking processes, with curiosity *per se* (e.g. Jepma et al., 2012). Given these facts we hypothesized that dACC neurons would track current level of information gap.

Here we made use of the curiosity tradeoff task that we developed previously (Blanchard et al., 2015). This task is based a version of the observing task designed for macaques (Bromberg-Martin & Hikosaka, 2009; Roper, 1999). On each trial, subjects choose between two gambles with different stakes and then wait 2.25 seconds until they are rewarded. One option provides information about the resolution of the gamble immediately; the other option maintains the mystery for the delay period. Monkeys are reliably information-seeking in this task, meaning they will sacrifice a small amount of water to obtain advance (Blanchard et al., 2015). We have proposed that this task satisfies an operational definition of curiosity (Wang & Hayden, 2019). Specifically, we believe that information-seeking choices in this task reflect a demand for information reflective of an information gap. Moreover, we believe that choice of an uninformative option leads to a state in which information is lacking and therefore maintains an information gap. In a previous study, we reported the responses of neurons in orbitofrontal cortex (OFC) during this task, although we did not examine either of these epochs (Blanchard et al., 2015). For the present study, we compared this dataset to a second dataset, collected at the same time as the first but not previously analyzed, recorded in dACC.

## RESULTS

### Behavior: macaques value advance information about gamble outcomes

We used a task we called the *curiosity tradeoff task* that we developed previously (Blanchard et al., 2015; see also Bromberg-Martin and Hikosaka, 2009, which motivated the design of our study).

#### Standard trials

(70% of trials): each gamble offers a 50% chance of a juice reward of varying amount (**Figure 1**). Regardless of choice, any reward is given 2.25 seconds later. Behavior of macaques in this task has been described in detail (Blanchard et al., 2015; Bromberg-Martin & Hikosaka, 2009; Bromberg-Martin, Matsumoto, & Hikosaka, 2010). Indeed, these two macaques were the same subjects used in our previous study and behavior here is, not surprisingly, nearly identical (Blanchard et al., 2015, **Figure 1C**). As in our previous study, both subjects preferred informative cues. Subjects B and H chose the gamble with higher stakes on 78.2% and 83.0% of trials (both are greater than chance, p<0.0001, binomial test). Subjects B and H chose the more informative option on 67.8% and 69.4% of trials respectively (both p<0.0001, binomial test). When the two options had equal stakes, both subjects preferred information (B: 78.8%, H: 78.1%). Indifference points (**Methods**) for the two subjects were 76 μl (B) and 51 μl (H). This indifference point identifies the subjective value of information.

**Figure 1.**
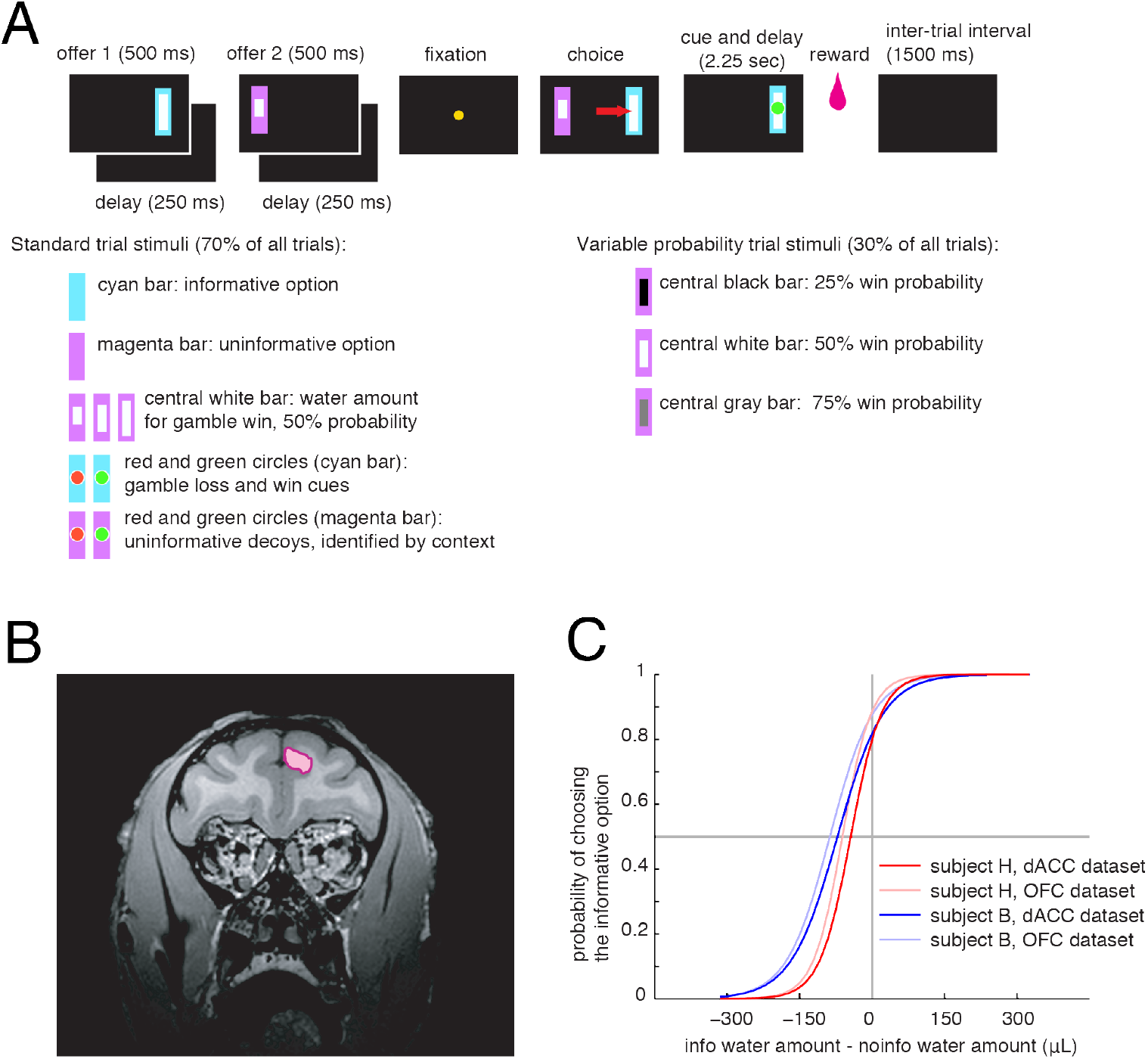
Task, anatomy, and basic behavior. **A**. Cartoon illustrating the structure of the task (above) and different possible stimuli (below). **B**. Coronal section of subject H showing the location of recording sites in dACC. **C**. Behavior of two subjects on standard trials in dACC/OFC datasets (darker/lighter colors). Likelihood of choosing informative option as a function of relative value between the two options. Leftward shift of curves indicates that both subjects preferred the informative option on standard trials.

#### Variable probability trials

(30% of trials): These trials were not used in our previous study and were introduced here as an additional control. On 30% of trials (randomly interleaved), subjects chose between two uninformative options that have the same stakes (225 uL juice). The probability was either 25%, 50%, or 70% and was the same for both offers on the same trial. On these trials, subjects chose the left and right option roughly equally (subject B: 55.1% left; subject H: 49.8% left). Any observed left/right bias did not depend on probability (regression of left choice against the three probability conditions, subject B: p=0.44, subject H: p=0.18).

### Enhanced pre-trial activity in dACC predicts information-seeking choices

We recorded responses of 151 single neurons in dACC (n=88 in subject B and n=63 in subject H). We collected an average 551 trials per cell, and a minimum of 500 trials. We reasoned that if demand for information reflects a drive state, it would have neuronal signatures before trial onset. We therefore considered the 500 ms period immediately preceding the presentation of the first offer. We divided all trials into two categories, (1) ones that were more information-seeking than average, (2) ones that were less information-seeking than average. These categories were defined in terms of the average subjective value the subject placed on information as inferred by the choice made during the task. Many trials could not be assigned to a category and were therefore excluded from this analysis (**Methods**).

For the example neuron shown in **Figure 2A**, pre-trial activity was higher on *relatively information-seeking* trials (p=0.004 Student’s t-test). Responses of 27.2% of neurons (n=41/151) differentiated the two trial types (this proportion is significant, p<0.001, binomial test, **Figure 2B**). Of these, 75.6% (n=31/41) showed enhanced firing (this proportion is significant, p=0.0015, binomial test). Responses of 26.4% of neurons (n=40/151) differentiated information-seeking trials relative to neutral trials (as determined by t-test, this proportion is significant, p<0.001, binomial test). Of these neurons, 70.0% (n=28/40) were enhanced (this proportion is significantly different from 0.5, p=0.0166, binomial test). Thus, increased pre-trial firing predicts information-seeking choices. Indeed, the average ensemble firing rate for all neurons (including non-significantly modulated ones) was 0.71 spikes/sec greater preceding information-seeking trials than neutral trials and 0.42 spikes/sec lower on information-averse trials than on neutral ones (both these differences are significant, p<0.001, t-test). These numbers represent a relatively high proportion (17.32% and 10.24%, respectively) of the baseline pre-trial firing rate (that is, 4.1 spikes/sec).

**Figure 2.**
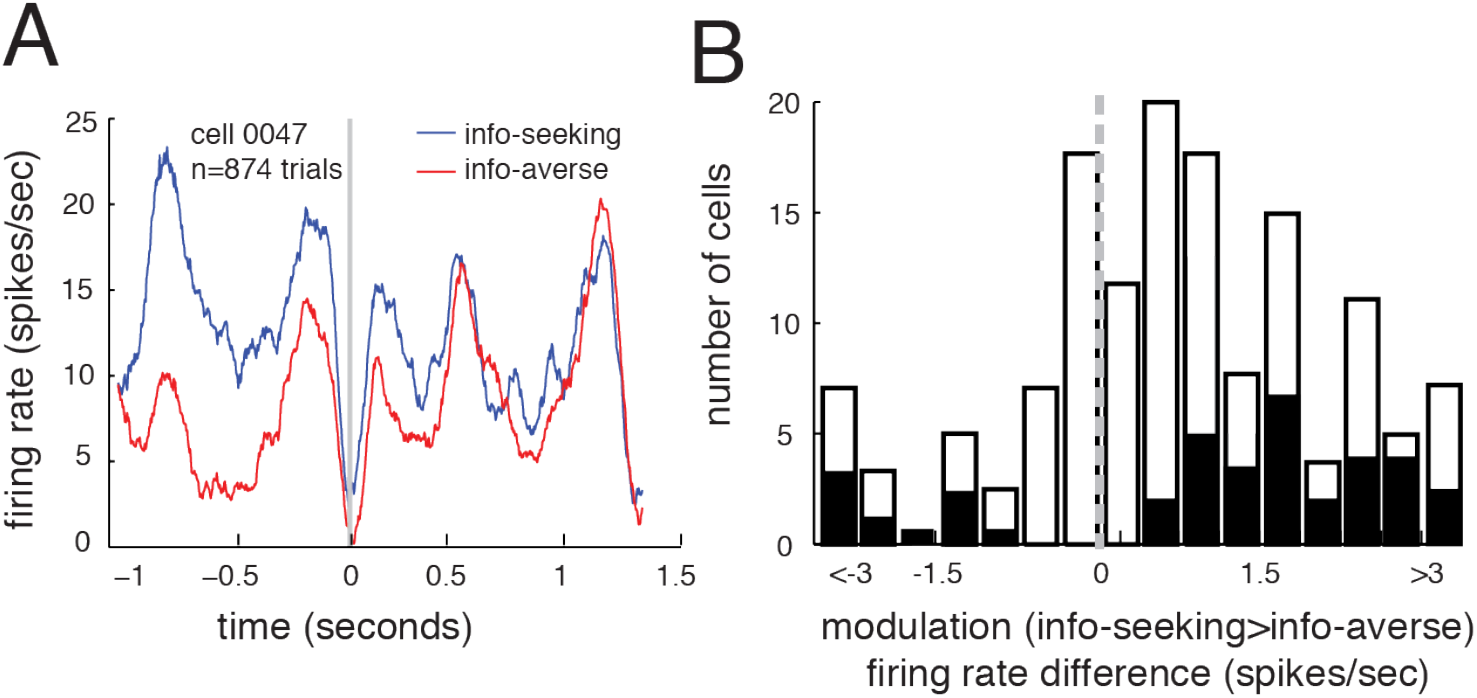
Pre-trial correlation between demand for information and firing rates in dACC neurons. **A**. Responses of an example neuron showing higher firing rates on info-seeking trials vs. info-averse trials (these categories are determined by average behavior, see main text). **B**. Histogram of pre-trial differences between info-seeking and info-averse trials. Neurons with individually significant effects are shown in black.

### Informational uncertainty tonically enhances firing rates in dACC

On trials in which the subject chose the no-info option (no-info trials), subjects proceeded to enter a state of temporally extended uncertainty. During this period, the subject did not know whether a reward would occur for 2.25 seconds. We next asked how neurons would respond to this sustained lack of uncertainty resolution. We reasoned that if uncertainty has no special implications, then the firing rate may resemble a weighted average of the firing rates associated with the two possible contrapositive outcomes (learning that a large/no reward is impending). On the other hand, if the status of lacking information in this task is somehow special, it may lead to a firing rate outside the range of the other two, and, in particular, systematic enhancement in firing across the long period the uncertainty is maintained.

For a typical neuron (**Figure 3A**), responses on no-info trials are enhanced (2.9 spikes/sec and 3.2 spikes/sec, p<0.01 in both cases, Student’s t-test). In our entire sample, firing rate on no-info trials was different from the average firing rate on both types of info trials in a substantial number of neurons (46.3%, n=70/151, p<0.001, binomial test). This modulation appears to last the entire waiting period. We divided the 2.25 second waiting period into nine equal 250 ms time bins. In all nine bins, a significant proportion of cells encoded the variables for info vs. no info. Even the bin with the lowest proportion had 18%, n=28 cells, which is greater than chance (p<0.001, binomial test).

**Figure 3.**
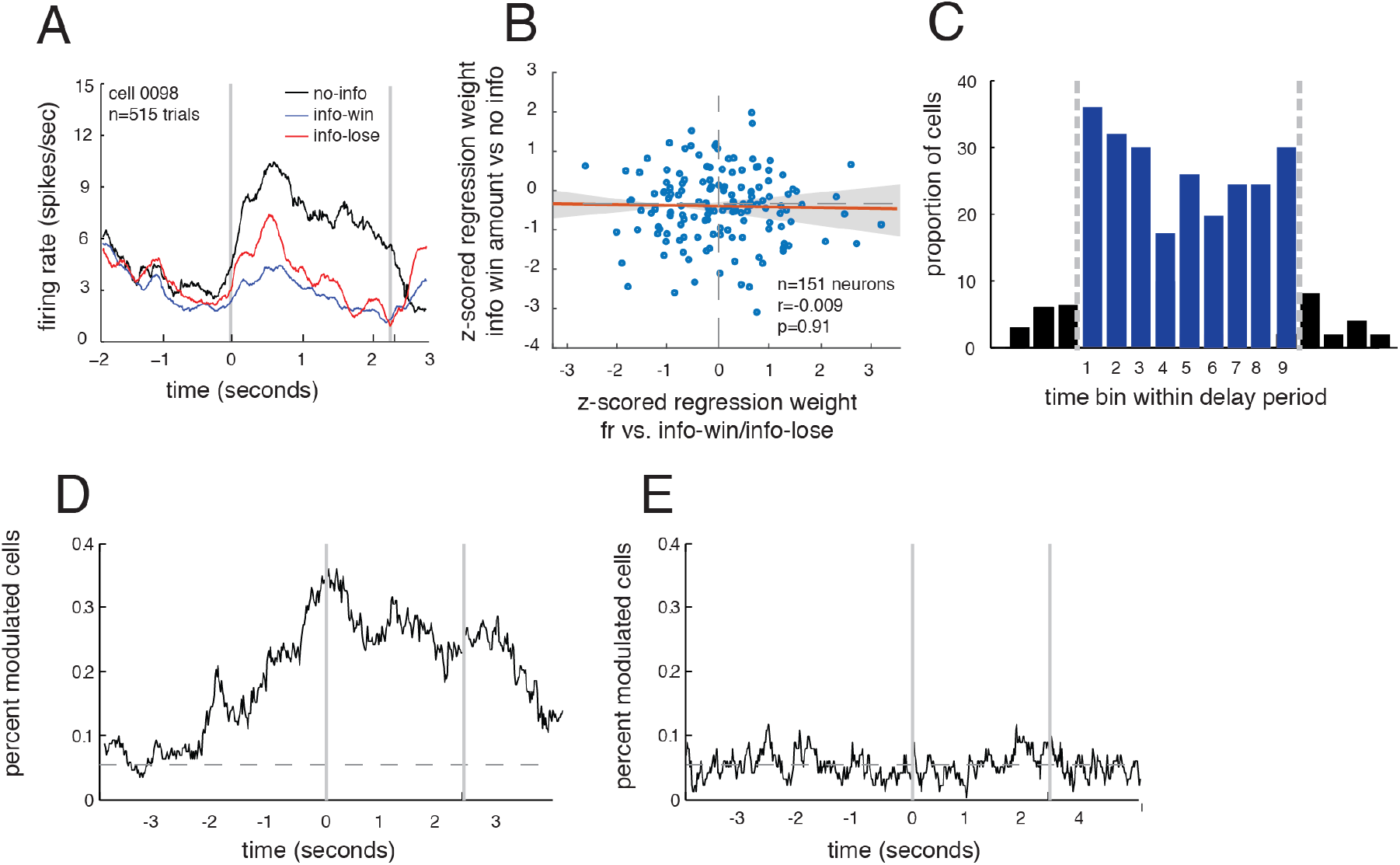
Delay period response modulation for no-info trials. Time 0 in A, D, and E indicates the start of delay period. **A**. Responses of an example neuron on trials in which no upcoming reward information is given during the delay (no-info trials, black) and trials in which this information is given (info trials, blue and red). Responses on no-info trials are systematically enhanced, a pattern that is common in the population. **B**. Scatter plot of regression weights for info vs. no info trials (y-axis) against info-win vs. info-lose (x-axis). These variables are not correlated, suggesting that codes for information gap and reward are unrelated. **C**. We divided data into nine time bins and found significant modulation in each one, suggesting, on no-info trials, the modulation is sustained across the delay period. **D**. Plot of proportion of cells significantly modulated by info vs. no-info status, using a sliding 500 ms window. Horizontal dashed line indicates chance level (i.e., 5%). **E**. Plot of proportion of cells significantly modulated by the win- and lose-related cues on no-info trials (when they are non-predictive). We see no measurable effect.

We next asked whether neurons that showed enhancement in one of these epochs were more likely to be the ones that showed enhancement in another. (That is, whether these effects reflect a sustained enhancement in some neurons, or periodic short bursting in more neurons). We reasoned that if the same set of cells was involved in signaling information from one bin to another, then we would see a positive correlation in their unsigned regression weights (i.e. absolute value of regression weights, see Azab & Hayden, 2017 for details). For every pair of bins (n=72 comparisons, i.e. 9 time bins x 8 other time bins), the cells involved were more overlapping than chance (correlation was significant, average r=0.29, p<0.05 in all individual cases).

We next considered the average effect of informational status on aggregate (grand average) firing rate. We found that the average firing rate on all no-info trials (8.22 spikes/sec) for all neurons (including non-significantly modulated ones) was greater than on info-trials (5.97 spikes/sec; this difference is significant, p<0.001, Student’s t-test). The population of significantly modulated cells was positively biased, meaning more individual neurons showed an increase in firing than showed a decrease (74.3%, n=52/70, p<0.001, binomial test).

Note that this average positive deflection is unlikely to reflect a sustained version of the bias the predicted information-seeking choices (see previous section). That bias led to greater firing before info trials, whereas the delay period modulation showed the reverse pattern. Thus, any firing rate hysteresis would presumably have reduced our measured effects, not spurred a false positive.

### Delay period enhancement is greater on high information-demand trials

In a previous study using this task, we found that the value of information (willingness to pay) rises with stakes of the chosen option (Blanchard et al., 2015). These results indicate that demand for information is higher on higher stakes trials (i.e. trials on which the subjects are in suspense about a higher valued gamble). Overall, responses of 21.9% of cells (n=33/151) were modulated by the stakes during the no-info delay period; the majority (72.7%, n=24/33) showed an enhancement (this bias is significant, p=0.0135, binomial test). Indeed, the average firing of the population was greater in the top stakes tercile than in the bottom stakes tercile (difference in the entire population, 1.32 spikes/sec, p<0.001, Student’s t test). Nonetheless, the firing rate in the bottom tercile was greater than responses in either info-win or info-lose conditions (difference in the entire population, 1.91 spikes/sec, p<0.01, Student’s t test).

### Tonic firing rates in dACC encode upcoming reward information

Enhancement on no-info trials may be a consequence of reward encoding. For example, perhaps it is unpleasant to wait in suspense, or, conversely, it may be pleasant to wait in anticipation. We thus leveraged our ability to perform within-task characterization of reward sensitivity for each neuron. **Figure 4A** shows the choice-aligned responses of an example neuron separated by trial type, no-info (black), info-win (blue), and info-lose (red). The format is the same as in **Figure 3A**. For this neuron, responses following info-win trials were tonically higher than responses following info-loss trials (red vs. blue line, average difference, 3.6 spikes/sec, p<0.01, t-test). This pattern was typical of neurons in the sample (**Figure 4B**). Tonic changes in firing rate across the epoch were observed in 41% (n=61/151) of neurons depending on the win-loss status of the trial. This bias did not show a directionality; 47.5% (n=29/61) showed an enhancement; the bias is not significant (p=0.80, binomial test).

**Figure 4.**
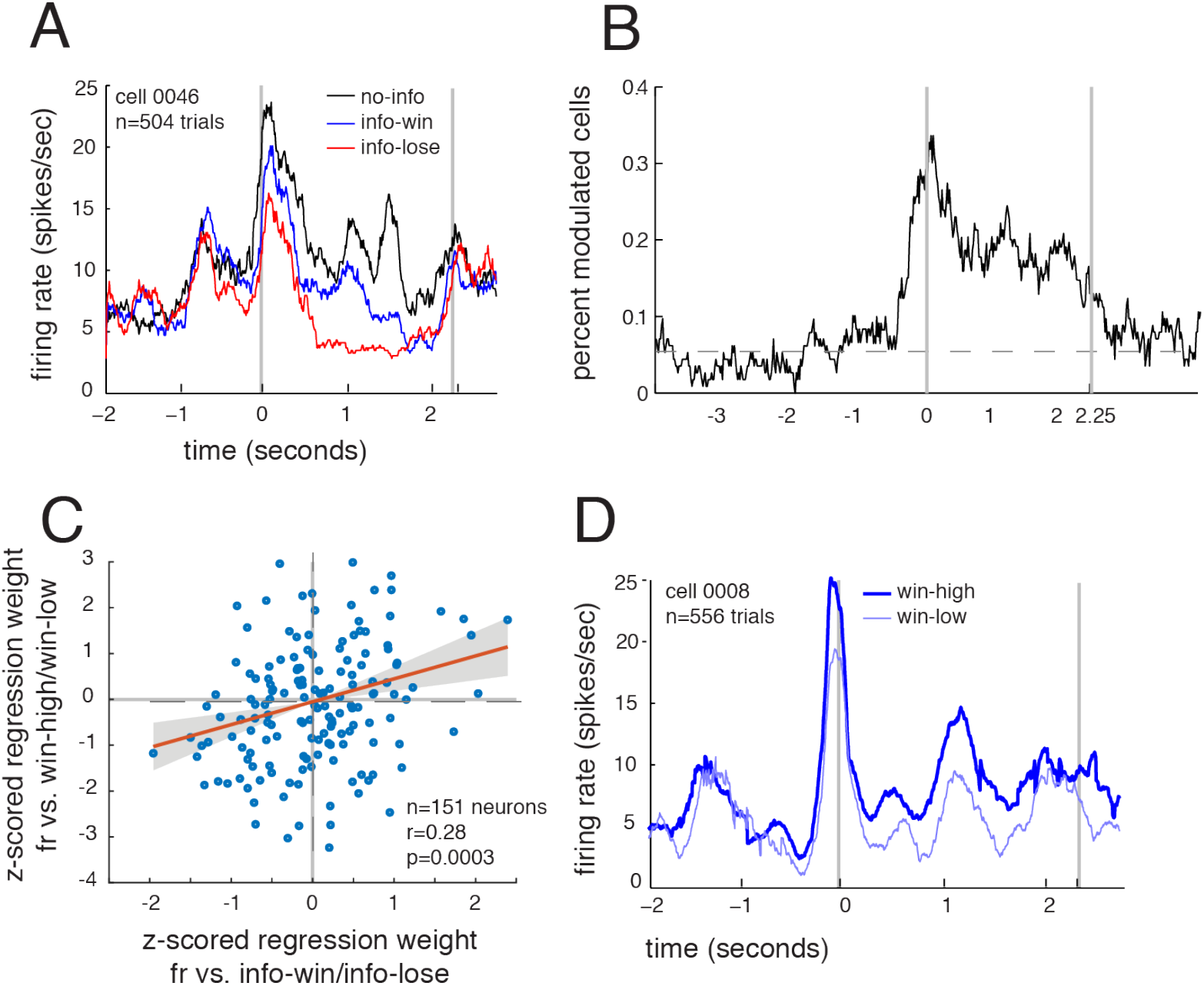
Neuronal encoding of upcoming rewards. **A**. Responses of an example neuron during the delay period (starting at time 0 on the graph) on info-win (blue) and info-lose (red) trials. Info-win and info-lose trials are significantly different throughout the course of the delay (0 to 2.25 seconds). Firing rates on no-info trials (black) are also shown, for reference. **B**. Proportion of cells whose responses significantly modulated by the difference between info-win and info-lose using a sliding 500 ms window. Horizontal dashed line indicates chance level (i.e., 5%). **C**. Scatter plot showing regression weights for info-win-high vs. info-win-low (y-axis) against info-win / info-lose (x-axis). The positive correlation indicates that dACC neurons use correlated codes for the two value variables. **D**. Responses of an example cell to info-win trials when the stakes are high (thick line) and low (thin line).

Neurons did not just encode win vs. loss. They also encoded specific reward volume anticipated. For the neurons in **Figure 4D**, the average firing rate was higher on info-win trials with larger than average rewards (thick line) than with smaller than average rewards (thin line, difference, 2.1 spikes/sec, p=0.009). On info-win trials, responses of 29.8% of neurons (n=45/151) encoded the stakes of the anticipated reward (regression of firing rate against size of anticipated reward). This bias was also not directional (19 positive and 26 negative, p=0.37, binomial test). The neural coding pattern, namely strength and direction, used by dACC neurons for the win-loss bias was closely correlated with that the reward volume effect, suggesting that this effect reflects a generic reward encoding (correlation of tuning indices for the two dimensions, r=0.31, p<0.001). This correlation indicates that, within dACC, there is a general code for anticipated reward - that is win vs. loss uses the same coding format as amount won on win trials. In any case, this result suggests that the lack of correlation between codes for information gap and for reward vs. no reward (see above and **Figure 2A**) is not simply and artifact of noise. (And indeed, that correlation, r=-0.02 is significantly lower than this one, Fisher r-to-z, z=-2.97, two-tailed p=0.003, see below).

### Controlling for confounds with reward and arousal in dACC

Subjects’ willingness to pay for information suggests it has an intrinsic value. We therefore wondered whether the tonic firing rate enhancement associated with lack of information is an artifactual consequence of reward or reward anticipation coding. We reasoned that if the information gap induced enhancement were an artifactual consequence of reward or reward anticipation coding, then we should see the neural coding pattern for information gap and reward related variables to be significantly similar. Otherwise, information gap evokes a differentiable pattern than do reward related variables in dACC. To test this idea, we therefore computed an *info-gap coefficient* (the linear term of the regression coefficient for firing rate against no-info vs info) and two reward indices for each neuron, one related to info-win vs. info-lose (*win-lose coefficient*) and one related to the size of the anticipated reward on the info-win trials (*win-amount coefficient*, see **Methods**).

The correlation between the info-gap coefficient and the two reward indices was not significantly different from zero in either case (win-lose coefficient: r=-0.02, p=0.59; win amount coefficient: r=-0.033, p=0.36). Because this lack of effect is difficult to interpret - it may reflect noise - we next estimated sample noise using a previously developed cross-validation technique, Blanchard et al., 2015). The correlations we observed within sample for info-gap coefficient with the two reward coefficients were both significantly greater than zero (r=0.67 and r=0.38, respectively, p<0.01 in both cases). They were also significantly greater than the observed correlations (differences were p<0.001 in both cases, bootstrap test), indicating that noise was not a limiting factor and indicating that our observed correlation was significantly less than the value we would have observed had the true correlation been 1.0. These results suggest that information gap and arousal (both reward-related coefficients) evoke unrelated neural response patterns and thus the effect of information gap cannot be simply explained away by arousal.

It is also worth noting that the modulation observed on no-info trials does not appear to reflect the low level features of the stimuli; on info and no-info trials, the same two cues were presented, but they had either reward-predictive or reward-irrelevant meaning, depending on context (**Figure 3E**). On no-info trials, dACC neurons did not encode the color of the decoy cue (5.3% of cells did so, n=8/151, p=0.85).

To gain additional perspective on the potential confound with arousal, we included a new trial type. On variable probability trials (30% of all trials), subjects chose between identical offers. These trials had either 25%, 50%, and 75% stakes and a medium reward. Responses of two example neurons are shown in **Figures 5A and 5B**. These neurons showed greater firing on 50% trials than on the other two trial types. Overall, 52.3% of neurons (n=79/151) showed a significant difference for the conditions (ANOVA test on individual neurons).

**Figure 5.**
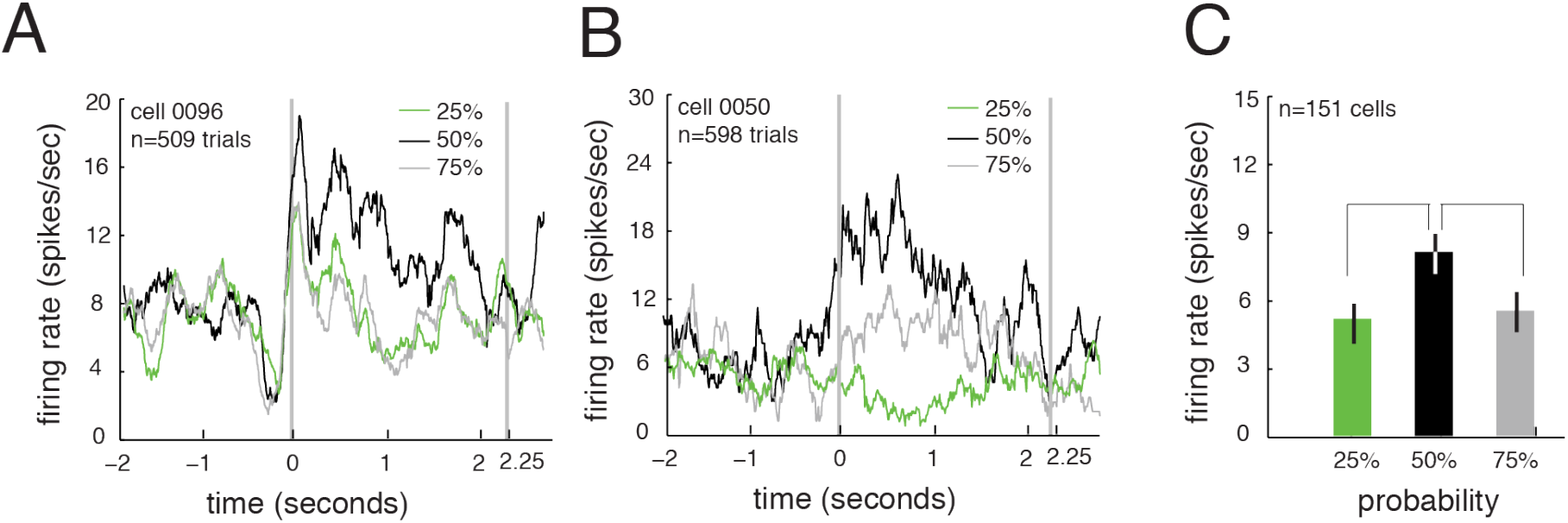
Data from variable probability trials. **A. and B**. Responses of two example neurons on *variable probability trials*. For both neurons, responses were greater on 50% trials than they were on 25% and 75% trials, suggesting the neurons is more concerned with entropy than it is with expected value. Time 0 refers to the start of the delay period. **C**. Grand average of responses for the population on variability probability trials. Responses of the ensemble are greater on 50% than on either 25% or 75% trials.

The example neurons are typical - we found that on these trials, neurons differentiated 25% from 50% (difference for all neurons: 3.46 spikes/sec, p<0.001), and 50% from 75% (difference for all neurons: 2.68 spikes/sec, p<0.001), although they did not differentiate 25% from 75% (difference: 0.39 spikes/sec, p=0.34). Note that these analyses reflect control for multiple comparisons. This pattern suggests that neurons encode entropy (sometimes called uncertainty), rather than expected value. In other words, the most parsimonious explanation of the factors driving neural responses is “amount of information available.” To formally test this idea, we compared linear and quadratic models; we found that the quadratic model fit better in more of the condition-selective neurons (n=38/79 for quadratic and 6/79 for linear fit, see **Methods** and Burnham & Anderson, 2010).

### Variations in dACC firing rate predict likelihood of changing strategy in response to outcomes

We wondered if the firing rate enhancement we saw correlates with readiness to learn. We have previously investigated the effects of risky outcomes on behavioral adjustments in some detail (Hayden et al., 2009; Hayden et al., 2011). For present purposes, the key idea is that switching – whether or not it is beneficial – is driven by attention to recent outcomes, so that variability in propensity to switch reflects variability in receptivity to recent outcomes.

Here, we find that following wins, subjects are more likely to choose the same side (left vs right). Specifically, relative to losses, on wins, subject B showed a 9.6% increased likelihood of repeating the rewarded side and subject H showed a 10.0% increase (these numbers, while small, are significantly greater than 0, p<0.001, binomial test, **Figure 6A**). These effects are observable as far out as 4 trials later. Gamble wins changed preference at a statistically significant level for subject B (3.5% increase, p=0.0288) and for subject H (3.8% increase, p=0.0446).

**Figure 6.**
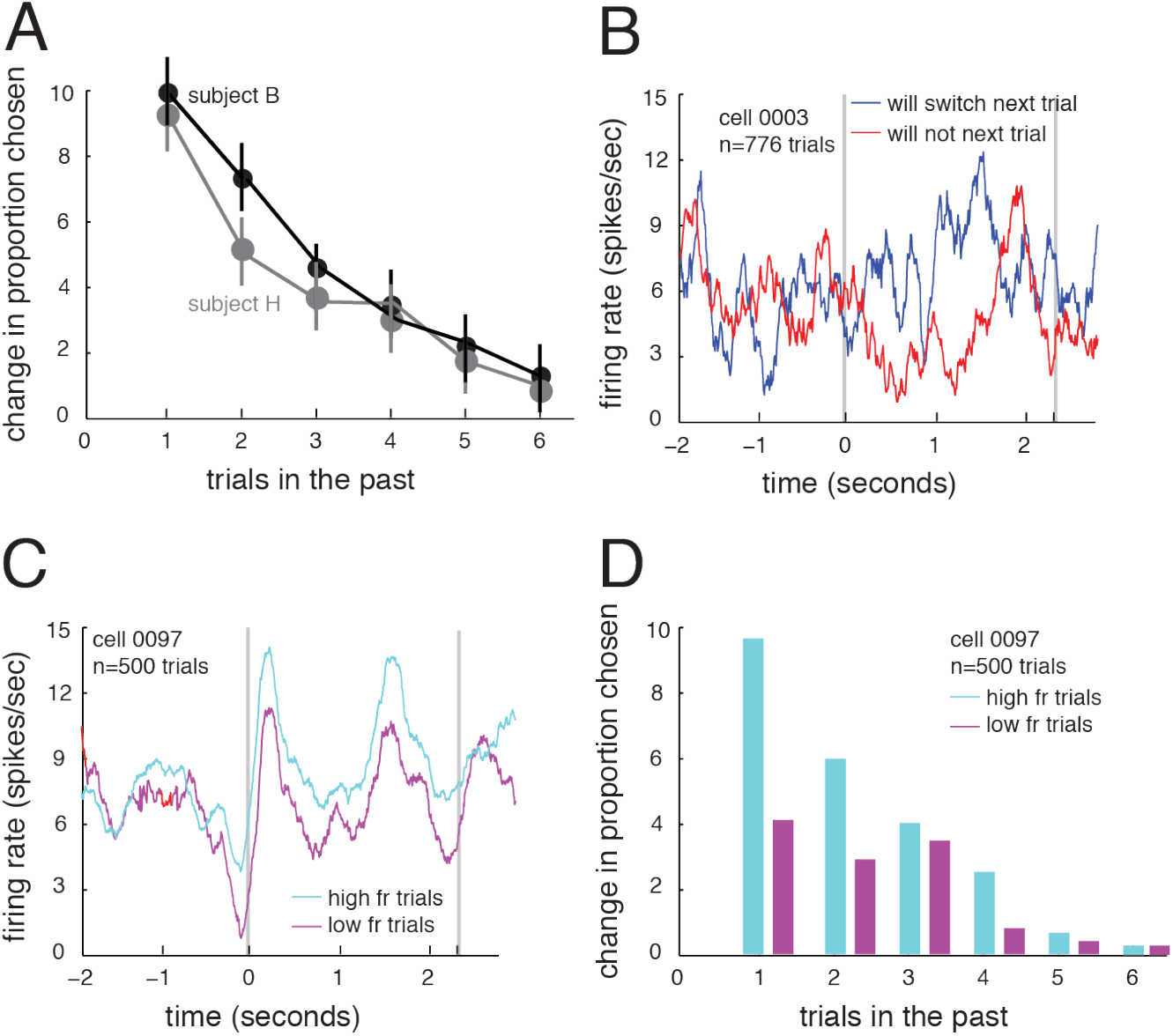
Data related to adjustment and likelihood of changing strategy. 0 point on X axis in A-C reflect the start of delay period. **A**. Subjects switch sides more often following gambling losses than gambling wins; this effect persists 3-4 trials. **B**. Plot of firing rate of an example cell showing different firing rates on on-info trials (i.e. controlling for information status and reward status) separated by whether the subject will switch on the next trial. **C**. Plot of an example cell in high and low firing rate trials (note that this effect, while significant, is a consequence of our analysis). Time zero indicates start of delay period. **D**. On higher firing rate trials for the neuron shown in panel B, subjects are more likely to adjust behavior on subsequent trials.

We next asked how these trial-to-trial adjustment effects correspond to variations in firing rate. **Figure 6B** shows the delay period firing rate of an example neuron on no info trials. This neuron showed enhanced firing rate for info-gap and this firing rate predicted choice switch on the next trial. **Figure 6C** shows the responses of an example neuron on no-info trials, separated into higher and lower than average firing rates, after regressing out stakes. For this neuron, responses were 1.81 spikes/sec higher on higher firing rate trials (p=0.019, t-test; note that this difference is pre-ordained by the analysis). **Figure 6D** then shows the adjustment pattern for this session on both trial types. This overall pattern was also observed in the population. Specifically, we performed a linear regression of firing rate in the window against side switch (binary variable, 1 for switch, - 1 for no-switch), including additional factors for stakes and past win/lose. Responses of 22.5% (n=34/151) of neurons show a correlation with switching, after regressing out other variables; 76.4% are positive (n=26/34, this proportion is significant, p=0.0029, binomial test). We also performed a linear regression of firing rate in the window against strategy switch (binary variable), including additional factors for stakes and past win/lose. We find that responses of 12.5% of neurons (n=19/151) show a significant correlation with switching, and that 15% are positive (n=15/19, p=0.0192).

### Lack of corresponding effects in OFC

We collected complementary results in a study of the OFC (those data are summarized in Blanchard et al., 2015). In our previous manuscript reporting on that dataset, our analyses focused on responses to offers, whereas here we consider pre-trial and delay period effects. Here, we report new results focusing on the pre-trial and delay period effects in the OFC dataset.

Overall, OFC appears very weakly involved in the aspects of the task that strongly drive dACC responses. In OFC, variability in firing rates pre-trial did not predict information-seeking decisions. Specifically, 7.96% of neurons (n=9/113 neurons) showed a firing rate correlation with upcoming choice (not significant, p=0.1877, binomial test). This proportion is significantly smaller than the proportion observed in dACC (i.e. 25.8%, p<0.001, binomial test). The average firing rate before information-seeking trials was not different than the average firing rate before information-averse trials (difference: 0.11 spikes/sec, p=0.85, t-test). Uncertainty about upcoming rewards did enhance delay activity in OFC in a significant proportion of neurons, although the proportion was close to threshold (9.7% of cells, n=11/113, p=0.0293, binomial test). The effect was visible as an increase in firing as in OFC, although the effect is not significant (difference: 0.44 spike/sec, p=0.33), and is significantly lower than the difference in dACC (p<0.001, Student’s t-test).

Finally, variation in firing rate in OFC did not predict adjustments in behavior. Specifically, we observed this correlation in 2.65% of cells (n=3/113). This proportion is not significant (p=0.29, binomial test) and is significantly lower than the proportion observed in dACC (p<0.01, binomial test). These results together suggest that OFC does not strongly predict information seeking behavior or strategy adjustment after the resolution of epistemic uncertainty.

## DISCUSSION

Curiosity, a drive for non-instrumental information, clearly has multiple possible causes. Here, we asked whether those causes can include mitigating the costs associated with uncertainty. Specifically, we reasoned that remaining in a state of suspense may be aversive in part because it carries some metabolic costs. To test this hypothesis, we examined the responses of single dACC neurons during an information tradeoff task (Blanchard et al., 2015). We find that tonically enhanced firing rates in dACC predict information-seeking on a trial-by-trial basis, a potential neuronal correlate of the that is a hypothesized driver of curiosity. Choice of uninformative options leads to a sustained tonic enhancement in firing that persists until the information is provided. Variability in this enhancement predicts demand for information and sensitivity of the subject to outcome information (as assessed by adjustment behavior). These changes were not observed in OFC, suggesting that our putative enhancements in activity related to uncertainty are at least somewhat anatomically localized. These observations in turn endorse the idea that dACC serves in part to accumulate evidence for purposes of guiding action (Hayden et al., 2011; Hayden and Heilbronner, 2016; Hunt et al., 2018; Kolling et al., 2012).

Neurons in many prefrontal regions encode multiple task variables and, typically, neural correlates of task variables show a population-level balance of positive and negative responses. This overall balance likely reflects the fact that positive and negative deflections can both carry information and, because spiking is costly, there are metabolic benefits that accrue to a brain that can keep overall spiking levels low regardless of the situation. The putative correlates of information gap we introduce here, in contrast, are biased towards the positive direction. The bias towards the positive direction suggests the speculative possibility that encoding these variables imposes metabolic costs on dACC (Laughlin et al., 1998). These costs did not appear to be counteracted by savings in other task epochs or in at least one other brain region, the OFC. If the brain is efficient at managing its own energy budget, it will seek out situations that can reduce spiking. Thus, our results provide tentative evidence consistent with the hypothesis that demand for information in this task reflects a demand for energy efficiency.

Why would it be costly to do be in a state of suspense? One possibility is that, when there is information available to learn, the brain’s learning systems enter into a state of *eligibility*, that is, they have the ability to enter into multiple possible knowledge states. Perhaps these knowledge states are low-energy, but the metastable state in which multiple knowledge states are possible is higher energy. Another – not incompatible - possibility is that the brain must enter into a state of enhanced vigilance to monitor information and that the acquisition of that information allows the brain to reduce its vigilance and focus on other tasks. Both possible explanations – eligibility and vigilance have at least some support in the form of previous correlations with dACC activity.

We have proposed that this task satisfies an operational definition of curiosity (Wang & Hayden, 2018; Wang et al., 2018; Wang and Hayden, 2019). An influential theory of curiosity holds that the demand for information is often driven by an *information gap* (Golman & Loewenstein, 2015; Gottlieb et al., 2013; Kang et al., 2009; Loewenstein, 1994; Golman & Loewenstein, 2018; Loewenstein, 1994). That is, a decision-maker’s assignment of value to an informative option is caused in part by a disparity between *desired* and *actual* knowledge. In this view, lack of information is a drive state that can be sated by information. Consumption of information is rewarding and lack of it - when desired - is aversive or at least dystonic. The information gap is the central theoretical structure linking curiosity to psychology and ultimately to neuroscience (Golman & Loewenstein, 2018; Gottlieb & Oudeyer, 2018; Kidd & Hayden, 2016; Marvin & Shohamy, 2016; van Lieshout et al., 2018). Our results suggest that the information gap would have a specific and anatomically localized set of correlates, and that this set includes dACC and not OFC.

Our results have some bearing on debates about the ultimate nature of the dACC. The function of this region has long been linked to both monitoring and executive control, as well as to core economic functions (Heilbronner & Hayden, 2016; Morecraft & Van Hoesen, 1998; Shenhav et al., 2013). Our work is most directly associated with theories suggesting it is a general-purpose monitor and controller. For example, past work suggests that dACC monitors conflict, reward outcomes, and other factors that lead to control (Alexander & Brown, 2011; Azab & Hayden, 2018; Botvinick et al., 1999; Shenhav et al., 2013; Shenhav et al., 2017; Hillman & Bilkey, 2010; Widge et al., 2019). Our results, then, suggest a tentative link between executive control and information-seeking, one that has been generally under-appreciated in the curiosity literature. In particular, they suggest that curiosity may serve be part of a larger tradeoff that involves efficient allocation of cognitive resources.

Functional neuroanatomy – the identification of region-specific functions is an important goal of cognitive neuroscience. Some cognitive functions related to economic choice appear to be broadly distributed (Cisek & Kalaska, 2010; Hunt & Hayden, 2017; Vickery et al., 2011; Yoo & Hayden, 2018). Our work here, however, indicates that there is what appears to be a qualitative difference between OFC and dACC (Kennerley et al. 2011; Rudebeck et al., 2006; Hunt et al., 2018). Because we were only able to record in two regions it is unclear what the full meaning of this difference is - one possibility is that monitoring is a specialized cingulate function. Another possibility is that OFC is specialized. Indeed, it has been proposed that OFC encodes a cognitive map of the state space for the currently relevant task but is not directly involved in changing behavior (Schuck et al., 2016; Wikenheiser & Schoenbaum, 2016; Wilson et al., 2014). If so, then it would not be involved in driving the state change or in keeping track of the environmental variables for potential state update. Our data suggest that dACC is a strong candidate for these functions, and may thus play a complementary role to OFC in this process.

## MATERIALS AND METHODS

### Electrophysiological Techniques

Two male rhesus macaques (*Macaca mulatta*) served as subjects. All procedures were approved by the University Committee on Animal Resources at the University of Rochester and were designed and conducted in compliance with the Public Health Service’s Guide for the Care and Use of Animals. In this manuscript, we discuss two related datasets, one from dACC (the focal dataset) and one from OFC (the comparator dataset). The same subjects were used for both studies; OFC data were collected first and the dACC dataset was collected soon afterwards using the same recording methods.

A Cilux recording chamber (Crist Instruments) was placed over the prefrontal cortex, overlying both area 24 of dACC (as defined in Heilbronner and Hayden, 2016a). This is the same region used in our past studies, e.g. Hayden et al., 2011; Hayden et al., 2009. We also recorded in area 13 of OFC (**Figure 1B**; this is the same region used in these subjects in our past studies, for example Wang and Hayden, 2017 and Sleezer et al., 2016). Position was verified by magnetic resonance imaging with the aid of a Brainsight system (Rogue Research Inc.). Neuroimaging was performed at the Rochester Center for Brain Imaging, on a Siemens 3T MAGNETOM Trio Tim using 0.5 mm voxels.

Single electrodes (Frederick Haer & Co., impedance range 0.8 to 4 mohm) were lowered using a microdrive (NAN Instruments) until waveforms were isolated. Action potentials were isolated on a Plexon system (Plexon, Inc). Neurons were selected for study solely based on the quality of isolation. All collected neurons for which we managed to obtain at least 500 trials were analyzed. Eye position was sampled at 1,000 Hz by an infrared eye-monitoring camera system (SR Research). Stimuli were controlled by a computer running MATLAB (Mathworks) with Psychtoolbox and Eyelink Toolbox. A standard solenoid valve controlled the duration of juice delivery. The relationship between solenoid open time and juice volume was established and confirmed before, during, and after recording.

### Information tradeoff task

Two offers were presented in sequence on each trial. The first offer appeared for 500 ms, followed by a 250 ms blank period; a second option appeared for 500 ms followed by a 250 ms blank period. Every trial had one informative and one uninformative option. The order of presentation (informative vs. uninformative) and location of presentation (info-on-left vs. info-on-right) varied randomly by trial. The offered water amount varied randomly for each option (75 to 375 μL water in 15 μL increments). 70% of trials were <u>standard trials</u>; for the OFC dataset, 100% of trials were standard trials. The remaining trials were <u>variable probability</u> trials; these were interleaved randomly with standard trials.

Each offer was represented by a rectangle 300 pixels tall and 80 pixels wide (11.35 degrees of visual angle tall and 4.08 degrees wide). On standard trials, all options offered a 50% probability of gamble win, to be delivered 2.25 seconds after the choice. Informative gambles (cyan rectangle) indicated that the subject would see a 100% valid cue immediately after choice indicating whether the gamble was won or lost.

Uninformative gambles (magenta rectangle) indicated that a random cue would appear immediately after choice. Valid and invalid cues were physically identical (green and red circles inscribed on the chosen rectangle). Each offer contained an inner white rectangle. The height of this rectangle linearly scaled with the water amount to be gained in the case of a gamble win. Offers were separated from the fixation point by 550 pixels (27.53 degrees). Subjects were free to fixate upon the offers (and almost always did so). After the offers, a central fixation spot appeared. Following 100 ms fixation, both offers reappeared simultaneously and the animal chose one by shifting gaze to it. Then the 2.25 s delay began, and the cue was immediately displayed. Any reward was delivered after this delay. All trials were followed by a 750 ms inter-trial interval (ITI) with a blank screen. Previous training history for these subjects at the time of recording included a full session (several months) with this task, two types of foraging tasks (Blanchard & Hayden, 2014; Hayden et al., 2011), three gambling/choice tasks (Farashahi et al., 2018; Heilbronner & Hayden, 2016b; Pirrone et al., 2018), and an attentional task (similar to the one used in Hayden & Gallant, 2013).

### Indifference point

We identified when subjects chose informative and non-informative options with equal probability (50%-50%) and then calculated the difference in stakes (as in water amount) between the two options. We found that non-informative would have to have larger stakes than informative ones and this number is 76 μl for subject B and 51 μl for subject H. Therefore, the information equates to 76 μl of juice reward for subject B and 51 μl for subject H.

### Identifying information-seeking and information-averse trials

For the pre-trial analysis, we divided all trials into three categories, (1) ones that were more information-seeking than average (information-seeking trials), (2) ones that were less information-seeking than average (information-averse trials), and (3) ones for which we could not assign information-seeking with any confidence (neutral trials). First, we computed an *equivalent value* for the uninformative option by adding a session-wide average value of information for that subject (i.e. 76 μl for subject B and 51 μl for subject H). In effect, this means we computed the average information-seekingness of the session and then divided trials into ones that were more or less information-seeking than would be predicted given the average. Trials were placed into the first category if the subject chose the informative option and its value was less than the equivalent value of the uninformative option. Trials were placed into the second category if the subject chose the uninformative option and its equivalent value was less than the value of the informative option. Note that in many trials, the choice did not provide information germane to this question, and these were place into a third class. For example, if the informative option had a value greater than that of the uninformative one, the subject’s choice would not be classifiable.

### Model comparison

We used AIC weights to conduct model comparison and select the better fitting model.

For model comparison, AIC weights were calculated as following:

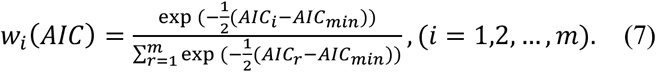

W_*i*_ is the probability of a model M_*i*_ being the one, among all *m* candidate models that is closest to the true data-generating model (Burnham & Anderson, 2010).

## Acknowledgements

We thank Tommy Blanchard for help in designing the task, and Marc Mancarella and Meghan Castagno for assistance in data collection. We appreciate close help from Ethan Bromberg-Martin for help in designing the task, and developing some of the analysis approaches we used here. This research was supported by an NIH R01(DA038106) to BYH.

## REFERENCES

Alexander, W. H., & Brown, J. W. (2011). Medial prefrontal cortex as an action-outcome predictor. Nature Neuroscience, 14(10), 1338–1344. http://doi.org/10.1038/nn.2921

Azab, H., & Hayden, B. Y. (2017). Correlates of decisional dynamics in the dorsal anterior cingulate cortex. PLoS Biology, 15(11), e2003091.

Azab, H., & Hayden, B. Y. (2018). Correlates of economic decisions in the dorsal and subgenual anterior cingulate cortices. The European Journal of Neuroscience, 47(8), 979–993.

Blanchard, T. C., & Hayden, B. Y. (2014). Neurons in Dorsal Anterior Cingulate Cortex Signal Postdecisional Variables in a Foraging Task, 34(2), 646–655. http://doi.org/10.1523/JNEUROSCI.3151-13.2014

Blanchard, T. C., Hayden, B. Y., & Bromberg-Martin, E. S. (2015). Orbitofrontal cortex uses distinct codes for different choice attributes in decisions motivated by curiosity. Neuron, 85(3), 602–614. http://doi.org/10.1016/j.neuron.2014.12.050

Botvinick, M., Nystrom, L. E., Fissell, K., Carter, C. S., & Cohen, J. D. (1999). Conflict monitoring versus selection-for-action in anterior cingulate cortex. Nature Publishing Group, 402(6758), 179.

Bromberg-Martin, E. S., & Hikosaka, O. (2009). Midbrain Dopamine Neurons Signal Preference for Advance Information about Upcoming Rewards. Neuron, 63(1), 119–126. http://doi.org/10.1016/j.neuron.2009.06.009

Bromberg-Martin, E. S., Matsumoto, M., & Hikosaka, O. (2010). Dopamine in motivational control: rewarding, aversive, and alerting. Neuron, 68(5), 815–834. http://doi.org/10.1016/j.neuron.2010.11.022

Cervera, R. L., Wang, M. Z., & Hayden, B. (2020). Curiosity from the Perspective of Systems Neuroscience. PsychArxiv.

Cisek, P., & Kalaska, J. F. (2010). Neural mechanisms for interacting with a world full of action choices. Annual Review of Neuroscience, 33, 269–298.

David, S. V., & Hayden, B. Y. (2012). Neurotree: A collaborative, graphical database of the academic genealogy of neuroscience. PloS one, 7(10).

Farashahi S, Azab H, Hayden B, Soltani A. (2018) On the flexibility of basic risk attitudes in monkeys. J. Neurosci. 38, 4383 – 4398. (doi:10.1523/jneurosci.2260-17.2018)

Golman, R., & Loewenstein, G. (2015). Curiosity, Information Gaps, and the Utility of Knowledge. SSRN Electronic Journal. http://doi.org/10.2139/ssrn.2149362

Golman, R., & Loewenstein, G. (2018). Information gaps: A theory of preferences regarding the presence and absence of information. Decision, 5(3), 143.

Gottlieb, J., & Oudeyer, P.-Y. (2018). Towards a neuroscience of active sampling and curiosity. Nature Reviews. Neuroscience, 1.

Gottlieb, J., Oudeyer, P.-Y., Lopes, M., & Baranes, A. (2013). Information-seeking, curiosity, and attention: computational and neural mechanisms. Trends in Cognitive Sciences, 17(11), 585–593. http://doi.org/10.1016/j.tics.2013.09.001

Hayden, B. Y., & Gallant, J. L. (2013). Working memory and decision processes in visual area v4. Frontiers in Neuroscience, 7, 18. http://doi.org/10.3389/fnins.2013.00018

Hayden, B. Y., Pearson, J. M., & Platt, M. L. (2011). Neuronal basis of sequential foraging decisions in a patchy environment. Nature neuroscience, 14(7), 933.

Hayden, B. Y., Heilbronner, S. R., Pearson, J. M., & Platt, M. L. (2011). Surprise signals in anterior cingulate cortex: neuronal encoding of unsigned reward prediction errors driving adjustment in behavior. Journal of Neuroscience, 31(11), 4178–4187.

Hayden, B. Y., Pearson, J. M., & Platt, M. L. (2009). Fictive reward signals in the anterior cingulate cortex. science, 324(5929), 948–950.

Heilbronner, S. R. (2017). Modeling risky decision-making in nonhuman animals: shared core features. Current opinion in behavioral sciences, 16, 23–29.

Heilbronner, S. R., & Hayden, B. Y. (2016a). Dorsal Anterior Cingulate Cortex: A Bottom-Up View. Annual Review of Neuroscience, 39(1), annurev–neuro–070815–013952. http://doi.org/10.1146/annurev-neuro-070815-013952

Heilbronner, S. R., & Hayden, B. Y. (2016b). The description-experience gap in risky choice in nonhuman primates. Psychonomic Bulletin & Review, 23(2), 593–600. http://doi.org/10.3758/s13423-015-0924-2

Hillman, K. L., & Bilkey, D. K. (2010). Neurons in the rat anterior cingulate cortex dynamically encode cost–benefit in a spatial decision-making task. Journal of Neuroscience, 30(22), 7705–7713.

Hunt, L. T., & Hayden, B. Y. (2017). A distributed, hierarchical and recurrent framework for reward-based choice. Nature Reviews. Neuroscience, 18(3), 172.

Hunt, L. T., Malalasekera, W. N., de Berker, A. O., Miranda, B., Farmer, S. F., Behrens, T. E., & Kennerley, S. W. (2018). Triple dissociation of attention and decision computations across prefrontal cortex. Nature neuroscience, 21(10), 1471–1481.

Jepma, M., Verdonschot, R. G., van Steenbergen, H., Rombouts, S. A., & Nieuwenhuis, S. (2012). Neural mechanisms underlying the induction and relief of perceptual curiosity. Frontiers in Behavioral Neuroscience, 6, 5.

Kang, M. J., Hsu, M., Krajbich, I. M., Loewenstein, G., McClure, S. M., Wang, J. T.-Y., & Camerer, C. F. (2009). The wick in the candle of learning: Epistemic curiosity activates reward circuitry and enhances memory. Psychological Science, 20(8), 963–973.

Kennerley, S. W., Behrens, T. E. J., & Wallis, J. D. (2011). Double dissociation of value computations in orbitofrontal and anterior cingulate neurons. Nature Publishing Group, 14(12), 1581–1589. http://doi.org/10.1038/nn.2961

Kolling, N., Behrens, T. E., Mars, R. B., & Rushworth, M. F. (2012). Neural mechanisms of foraging. Science, 336(6077), 95–98.

Kidd, C., & Hayden, B. Y. (2016). The Psychology and Neuroscience of Curiosity. Neuron, 88(3), 449–460. http://doi.org/10.1016/j.neuron.2015.09.010

Laughlin, S. B., Van Steveninck, R. R. de R., & Anderson, J. C. (1998). The metabolic cost of neural information. Nature Neuroscience, 1(1), 36.

Loewenstein, G. (1994). The psychology of curiosity: A review and reinterpretation. Psychological Bulletin, 116(1), 75.

Marvin, C. B., & Shohamy, D. (2016). Curiosity and reward: Valence predicts choice and information prediction errors enhance learning. Journal of Experimental Psychology: General, 145(3), 266.

Morecraft, R. J., & Van Hoesen, G. W. (1998). Convergence of limbic input to the cingulate motor cortex in the rhesus monkey. Brain Research Bulletin, 45(2), 209–232.

Pirrone, A., Azab, H., Hayden, B. Y., Stafford, T., & Marshall, J. A. R. (2018). Evidence for the speed–value trade-off: Human and monkey decision making is magnitude sensitive. Decision, 5, 129–142. doi 10.1037/dec0000075

Roper, K. L. E. A. (1999). Observing Behavior in Pigeons: The Effect of Reinforcement Probability and Response Cost Using a Symmetrical Choice Procedure, 1–20.

Rudebeck, P. H., Buckley, M. J., Walton, M. E., & Rushworth, M. F. S. (2006). A role for the macaque anterior cingulate gyrus in social valuation. Science, 313(5791), 1310–1312. http://doi.org/10.1126/science.1128197

Schuck, N. W., Cai, M. B., Wilson, R. C., & Niv, Y. (2016). Human Orbitofrontal Cortex Represents a Cognitive Map of State Space. Neuron, 91(6), 1402–1412. http://doi.org/10.1016/j.neuron.2016.08.019

Shenhav A, Musslick S, Lieder F, et al. (2017). Toward a Rational and Mechanistic Account of Mental Effort. Annu Rev Neurosci. 40: 99–124.

Shenhav, A., Botvinick, M. M., & Cohen, J. D. (2013). The expected value of control: an integrative theory of anterior cingulate cortex function. Neuron, 79(2), 217–240.

Sleezer, B. J., Castagno, M. D., & Hayden, B. Y. (2016). Rule encoding in orbitofrontal cortex and striatum guides selection. Journal of Neuroscience, 36(44), 11223–11237.

Smith, E. H., Horga, G., Yates, M. J., Mikell, C. B., Banks, G. P., Pathak, Y. J., … Sheth, S. A. (2019). Widespread temporal coding of cognitive control in the human prefrontal cortex. Nature neuroscience, 1–9.

Strait, C. E., Blanchard, T. C., & Hayden, B. Y. (2014). Reward value comparison via mutual inhibition in ventromedial prefrontal cortex. Neuron, 82(6), 1357–1366. http://doi.org/10.1016/j.neuron.2014.04.032

van Lieshout, L. L., Vandenbroucke, A. R., Müller, N. C., Cools, R., & de Lange, F. P. (2018). Induction and relief of curiosity elicit parietal and frontal activity. Journal of Neuroscience, 38(10), 2579–2588.

Vickery, T. J., Chun, M. M., & Lee, D. (2011). Ubiquity and specificity of reinforcement signals throughout the human brain. Neuron, 72(1), 166–177.

Wang, M. Z., & Hayden, B. Y. (2019). Monkeys are curious about counterfactual outcomes. Cognition, 189, 1–10.

Wang, M. Z., & Hayden, B. Y. (2017). Reactivation of associative structure specific outcome responses during prospective evaluation in reward-based choices. Nature Communications, 8, 15821. http://doi.org/10.1038/ncomms15821

White, J. K., Bromberg-Martin, E. S., Heilbronner, S. R., Zhang, K., Pai, J., Haber, S. N., & Monosov, I. E. (2019). A neural network for information seeking. Nature communications, 10(1), 1–19.

Widge, A. S., Heilbronner, S. R., & Hayden, B. Y. (2019). Prefrontal cortex and cognitive control: new insights from human electrophysiology. F1000Research, 8.

Wikenheiser, A. M., & Schoenbaum, G. (2016). Over the river, through the woods: cognitive maps in the hippocampus and orbitofrontal cortex. Nature Reviews. Neuroscience, 17(8), 1–11. http://doi.org/10.1038/nrn.2016.56

Wilson, R. C., Takahashi, Y. K., Schoenbaum, G., & Niv, Y. (2014). Orbitofrontal cortex as a cognitive map of task space. Neuron, 81(2), 267–279. http://doi.org/10.1016/j.neuron.2013.11.005

Yoo, S. B. M., & Hayden, B. Y. (2018). Economic choice as an untangling of options into actions. Neuron, 99(3), 434–447.

